# Evaluation of the safety and insecticidal efficacy of ivermectin-treated bird feed formulations in different avian species

**DOI:** 10.64898/2025.12.18.695243

**Authors:** Michelle J. Savran, Kathryn Coffin, Claire M. Stewart, Chilinh Nguyen, Catalina Puska, Molly E. Ring, Anna-Sophia Leon, Preston Schweiner, Tilman Peters, Jenna C. Randall, Jenny Buczek, Paula Lado, Emily Gallichotte, Brady Clapsaddle, Christopher M. Barker, Gregory D. Ebel, Brian D. Foy

## Abstract

**Background:** West Nile virus (WNV) is maintained in an enzootic cycle between reservoir host birds and *Culex* (*Cx.*) spp. mosquitoes. This relationship presents a potential target for vector control strategies. Ivermectin (IVM), an endectocidal drug that selectively affects invertebrates while remaining safe at high concentrations in mammals and birds, can be delivered to *Cx. tarsalis* via blood meals from birds fed IVM-treated bird feed. In this study we evaluated the safety, efficacy, and utility of IVM-treated bird feed as a novel vector control strategy by assessing its impact on multiple bird species and mosquitoes.

**Methods:** Mosquitoes were collected during peak WNV transmission season in Northern Colorado and DNA extracted from blood meals to determine host species. Chickens, pigeons, zebra finches, and house sparrows were fed different formulations of IVM-treated bird feed and observed for clinical signs, and their sera were fed to *Cx. tarsalis* mosquitoes to evaluate mosquitocidal efficacy. Feeding rates and IVM serum concentrations in birds were analyzed by unpaired t-test and one-way ANOVA, and mosquito survivorship was analyzed by Kaplan-Meier curves and compared using paired log-rank tests. IVM serum concentration and mosquito survivorship were compared using Spearman correlation.

**Results:** Speciation analyses conducted on blood meals from *Cx. tarsalis* collected during peak WNV transmission season in Northern Colorado determined that they feed primarily on songbird species that commonly visit bird feeders, with house sparrows representing the most frequent blood meal host. In laboratory experiments using multiple formulations and doses of IVM, chickens, pigeons, zebra finches, and house sparrows ate comparable amounts of IVM-treated bird feed compared to untreated feed, had similar weight gain, and exhibited no clinical signs of toxicity. Both colony-reared and locally captured *Cx. tarsalis* showed significant mortality after feeding on sera from IVM-treated birds compared to controls.

**Conclusions:** These results suggest that targeting songbirds with IVM-treated bird feed should be safe for wildlife and may elicit high rates of IVM-induced mortality by reaching a large proportion of WNV vector mosquitoes via their proclivity for feeding on passerine birds.

## Background

West Nile virus (WNV) is the leading cause of locally acquired mosquito-borne disease in the United States, causing significant disease and death in humans and animals every year [1–3]. Since the introduction of WNV to the USA in 1999, over 56,000 WNV disease cases and over 2,700 WNV-related deaths have been reported to the Center for Disease Control (CDC), with the highest incidence occurring in the western US and Great Plains region [4–6]. Moreover, models estimate the actual number of infections to be closer to 7 million with approximately 1 million individuals developing disease [7]. While most infected individuals remain asymptomatic, about 20% develop febrile illness and 1 in every 150 febrile cases develop neuroinflammatory conditions that often result in long-term disability or death [8,9]. WNV has also impacted the health of North American bird populations since its emergence on the continent, inducing significant population declines in several species which then result in increased susceptibility to other natural and anthropogenic impacts [3]. Additionally, more than 28,000 equine WNV cases have been reported to the United States Department of Agriculture (USDA) with an average morality rate of 30-40% [10]. Unfortunately, development and marketing of a WNV vaccine for human use remains unlikely in the foreseeable future due to the associated costs, epidemiological factors, and difficulties in planning and conducting clinical trials to prove its efficacy, leaving insecticides as the most practical strategy to control the spread of WNV [11,12]. Insecticides primarily target the mosquito vector with biological and chemical interventions such as larvicidal bacteria, insect growth modulators, and adulticidal pyrethroid sprays [13–15]. Larvicides are critical for integrated mosquito management for their low cost and high target specificity, but several predominant WNV vector species have developed resistances to the mechanisms of some of these agents and can be difficult to target in their larval habitats [16–18]. Adulticides can successfully reduce adult WNV vectors in treated areas throughout the USA, but adulticide applications can be expensive to implement, and success can vary by local infrastructure, thus limiting this option to wealthier urban and suburban communities [19–21]. Additionally, while numerous studies have demonstrated a strong safety profile when using the recommended application rates of the most often used adulticides, other studies have linked specific adulticides with toxicity for key pollinators, as well as threatened and endangered species which has led to negative community perceptions and decreased utilization [22–25]. These drawbacks combined with predicted future increases in WNV disease risk indicate an urgent need for an alternative means of transmission control [26].

WNV is maintained in an enzootic cycle between passerine birds and ornithophilic *Culex* species mosquitoes. This ornithophilic blood feeding habit can be used to target birds with novel interventions designed to control the spread of arboviruses like WNV [27,28]. We have previously investigated the use of ivermectin (IVM)-treated bird feed to selectively target the main WNV bridge vector, *Cx. tarsalis*, in the western USA [29–31]. IVM is an endectocide that targets the nerve and muscle cells of invertebrates while maintaining a robust safety profile in vertebrates [32–35]. Field research and modeling studies completed by some of our group suggest that IVM control strategies can also be used to control the spread of certain mosquito-borne pathogens such as malaria [36,37]. IVM and other endectocides have been effectively employed for decades as both topical and feed-through treatments for many animals, from rodents to livestock, for tick, flea, and sand fly control [38,39]. Because WNV is primarily transmitted by older mosquitoes due to the long extrinsic incubation period relative to the mosquito’s lifespan, infected mosquitoes that consume blood meals from IVM-treated birds are expected to have a high probability of dying before they can transmit WNV to another host [40]. Our previous studies have shown that *Cx. tarsalis* survival is significantly reduced following direct or indirect blood meals from chickens (*Gallus gallus*), wild Eurasian collared doves (*Streptopelia decaocto*), and a wild common grackle (*Quiscalus quiscula*) fed a simple formulation of IVM powder mixed into chicken feed (200 mg/kg feed) compared to controls [29]. In 2016, we placed IVM powder feed and control feed feeders in Fort Collins, CO. We observed doves and songbirds visiting all stations and detected IVM in sera of 13/15 (87%) of birds trapped around IVM treated sites [29]. In 2019, we randomized eight flocks of chickens in Davis, CA and fed four flocks the same simple powder formulation for 72 days [30]. We observed delayed and non-significantly decreased numbers of WNV-positive *Cx. tarsalis* pools at IVM sites relative to controls in 2016 and significantly reduced WNV seroconversion rates in chickens from IVM-treated flocks in 2019.

While our previous results indicate simple formulations of IVM-treated feed could impact WNV transmission in the field, there are outstanding gaps to address before pursuing evaluation in the field. We have observed birds at experimental feeder stations previously, but we need to confirm mosquitoes will feed on bird species likely to visit the feeders. Additionally, the simple formulation we tested previously does not protect the drug from degradation and so must be replaced daily in the field, making it an impractical solution for use. We have also not tested safety and efficacy for extended periods in small passerine birds. In this study, we assessed blood meal host preferences among mosquitoes captured in Northern Colorado to determine which species we should target with medicated feed for WNV control. We also tested new IVM-treated bird feed formulations to determine ones that may work best in the field and simultaneously be safe for, and induce mosquitocidal effects in, multiple bird species.

## Materials and Methods

### Blood meal host analysis

Mosquitoes were collected and processed as described previously [41]. Briefly, CDC Miniature Light Trap Model 512 traps were set without light and baited with dry ice on volunteer properties throughout northern Colorado (**Additional file 1: Table S1**, host data listed in **Additional file 1: Table S2**). Trapped arthropods were brought back to CSU, frozen at -20°C, and engorged *Cx. tarsalis* mosquitoes were identified. Engorged *Cx. tarsalis* mosquitoes were individually separated into 2 mL microcentrifuge tubes containing one sterile glass plating bead, 250 µL tissue lysis buffer and 20 µL Proteinase K solution from the Mag-Bind® Blood & Tissue DNA HDQ 96 Kit (Qiagen, Valencia, CA). Mosquitoes were homogenized at 24 Hz for 1 minute, centrifuged at 13,000 × *g* for 5 minutes, and incubated at 55°C in a shaking incubator overnight. DNA was extracted from mosquito homogenates via the Mag-Bind® Blood & Tissue DNA HDQ 96 Kit using a KingFisher™ Flex Purification System (ThermoFisher Scientific), according to the manufacturer’s instructions. Blood meals in extracted DNA were identified by polymerase chain reaction amplification and sequencing of a fragment of either the vertebrate mitochondrial cytochrome c oxidase 1 (COI) [42] or cytochrome b (cytb) [43,44] genes, using the primer sets and thermocycling conditions described (**Additional file 1: Table S3**). Samples were submitted to Azenta Life Sciences for Sanger sequencing. The Barcode of Life Data System database was used to identify COI sequences (www.boldsystems.org), and GenBank was used to identify cytb sequences.

### Mosquito maintenance and bioassays

Colonized *Cx. tarsalis* (Kern National Wildlife Refuge strain) were reared in standard insectary conditions (28°C, 16:8 light cycle). Approximately 150 larvae were reared in roughly 11 liters of water and fed 2.5 grams of powdered Tetramin fish food daily until pupation. Adults were housed in groups of approximately 300 per cage and fed sugar and water ad libitum until separated for bioassays. Field-collected *Cx. tarsalis* were captured in CDC Miniature Light Trap Model 512 traps (BioQuip) used without light and baited with dry ice. These were set throughout Fort Collins, CO (**Additional file 1: Table S4**) and brought back to CSU for holding. Trapped arthropods were knocked down by cold exposure and *Cx. tarsalis* were identified, then separated into groups of 1:10 male:female and kept in standard insectary conditions (described above) for subsequent bioassays. For bioassays, mosquitoes were separated into groups of 50 females and 10 males, then starved of sugar and water for up to 18 hours. Laboratory mosquitoes were provided artificial membrane blood meals [45] consisting of 50% avian serum and 50% red blood cells from defibrinated chicken or calf blood (Colorado Serum Company, CO, USA) via glass feeders (Lillie Glass Blowers, GA, USA) kept warm by a circulating water bath set to 37°C. Wild-caught mosquitoes do not readily feed from artificial membranes, so these mosquitoes were fed directly on birds as described previously [29]. Following 30 minutes of feeding, mosquitoes were knocked down by cold exposure and fully engorged females separated and mortality recorded daily for seven days.

### Animals

Birds were provided with clean water and fed control or IVM-treated feed for three weeks (chickens, **Figure 2**) or seven days (all other birds, **Figures 3-6**). Feed consumption was measured daily (pigeons, zebra finches, house sparrows) and birds were weighed weekly (chickens), daily (pigeons), or every other day (house sparrows, zebra finches) to minimize stress. All birds were monitored daily for symptoms of toxicosis including but not limited to mydriasis, abnormal stool, stupor, ptosis, and ataxia. Birds received a behavioral score of possible toxicity: A) <u>Normal</u>, no toxicity observed; B) <u>Possible toxicity</u> – with the action to monitor three times daily for signs of continued toxicity and for animals in the same group to have similar symptoms; C) <u>Continued toxicity for >2 days</u>, with the action that if noted as B more for than two days, to euthanize immediately; D) <u>More than 2 birds per group develop continued toxicity</u>, with the action that if more than two birds per treatment group were noted as C, to stop experiment with that dose and euthanize all animals in the group. All blood samples were allowed to clot for 30 minutes at room temperature, then separated into serum and red blood cells by centrifugation at 2,000 × *g* for 10 minutes. Serum was transferred to a 1.7 mL microcentrifuge tube and stored at -80°C until used for bioassays or IVM quantitation.

#### Chickens

Four-week-old white leghorn chickens (*Gallus gallus*) were purchased from Northern Colorado Feeders, divided into groups, and housed outdoors in separate chicken coops. All chickens were provided clean water and chick starter crumble mix (mash) from Northern Colorado Feeders Supply for a period of three weeks, fed *ad libitum*. Birds in the treatment group received 200 mg of powder IVM formulation (Merck & Co., Inc., Kenilworth, NJ, USA) per kg of bird feed, mixed fresh daily. Blood was collected from each bird via jugular venipuncture once every two weeks. To compare mortality between field-collected and colonized *Cx. tarsalis* while controlling for variability in IVM serum concentration across time, chicken blood for artificial membrane feeds was drawn on the same day that field-collected mosquitoes were fed directly on the same chickens. Chickens were bled on an alternating schedule to evaluate mosquitocidal efficacy of serum over time without exceeding biweekly maximum of 1% body weight blood collection.

#### Pigeons

Feral pigeons (rock doves, *Columba livia*) were captured in Larimer County, CO and housed indoors in separate flight cages. All pigeons were provided clean water and either control seeds, untreated IVM millet or sunflower seeds, or Opadry® II (Colorcon Inc, Harleysville, PA, USA)-coated IVM millet (TDA Research Inc, Golden, CO, USA). Blood was collected from each pigeon via brachial venipuncture on day three and day seven of the diet.

#### Zebra finches

Zebra finches (*Taeniopygia guttata*) were purchased from a private breeder. Zebra finches were separated into groups (n=3 for 50 mg/kg IVM diet, n=8 for 100 mg/kg IVM diet) and provided clean water and millet diets, receiving either control millet, untreated IVM millet, or Opadry II-coated IVM millet for seven days. Blood was collected from each zebra finch after seven days of the IVM diet via intracardiac puncture under terminal isoflurane anesthesia.

#### House sparrows

Wild house sparrows (*Passer domesticus*) were captured in Wellington, CO (40°42’57.8”N 105°01’09.9”W), divided into groups (n=5 per each IVM diet, n=3 per control diet) and housed indoors in separate flight cages. All sparrows were provided clean water and millet diets, receiving either control millet, untreated IVM millet, or Opadry II-coated IVM millet for seven days. Blood was collected from each house sparrow at the conclusion of the diet study via intracardiac puncture under terminal isoflurane anesthesia. Following euthanasia, house sparrow liver tissue was fixed in 10% neutral buffered formalin, trimmed and embedded in paraffin, sectioned at 5 µm, and stained with hematoxylin and eosin. Histopathological examination of liver tissue was performed by an anatomic pathologist that was blinded to treatment groups. The following lesional scoring system was used and the summation of scores was compared across individuals and treatment groups, 1: minimal changes, 2: mild, 3: moderate, 4: marked, 5: severe for each category (lipid vacuolization, hepatic nuclear toxic change, hepatic necrosis, and inflammatory cell infiltrate).

### IVM extraction and quantification by HPLC tandem mass spectrometry

25 µL serum was treated with 100uL of 200ng/mL abamectin (ABM) (Sigma-Aldrich, St. Louis, MO, USA) prepared in methanol (Honeywell, Charlotte, NC, USA) and vortexed until homogenous. Samples were precipitated by freezing at -80°C for 1 hour and centrifuged at 30,000 × *g* at 4°C for 30 minutes. The supernatant layer was collected and evaporated under vacuum for approximately one hour. The extract was reconstituted in 25 µL methanol, vortexed, and sonicated for 5 minutes. The resuspension was centrifuged for 30,000 × *g* at 4°C for 10 minutes, then supernatant was transferred to an HPLC vial with screw cap. Analyses were performed using an Agilent 1290 Infinity II liquid chromatograph system coupled to an Agilent 6470 triple quadrupole mass spectrometer electrospray ionization (ESI) source. Nitrogen dry gas flow rate was 10 L/min and held at 325□C. Sheath gas flow rate was 11 L/min and held at 300□C. Nebulizer gas pressure was 38 psi. The fragmentor voltage was 150 V. In positive polarity the voltage capillary was 3500 V with a nozzle voltage of 0 V. In negative polarity, the voltage capillary was 3500 with a nozzle voltage of 2000 V. One quantifying and one qualifying MRM transition was monitored for both analyte and internal standard, each with a 150 V cycle time. For ivermectin, the precursor ion to product ion transitions were m/z 873.5 → 229.2 (quantifying, collision energy (CE) = 30 V) and m/z 567.5 (qualifying, CE = V). For abamectin, the precursor ion to product ion transitions were m/z 871.5 → 229.2 (quantifying, collision energy (CE) = 30 V) and m/z 565.5 (qualifying, CE = 30 V).

Reversed-phase separation of analytes was achieved on a Waters BEH C8 column (2.5mm x 100mm, 2.5 micron) using a gradient of 5mM ammonium acetate in water (mobile phase A) and 5mM ammonium acetate in 95% acetonitrile, 5%methanol (mobile phase B). The programmed gradient was as follows: hold 10% B from 0–1 minute, linear increase to 99% B from 1-7 minutes, hold at 99% until 12 minutes, linear decrease to 10% from 12-12.5 minutes and hold 10% B to 16.5 minutes. Chromatographic flow rate remained constant at 0.3 mL/min and column temperature remained at 40□C.

The linearity of the method was tested after elaboration of analytical calibration curves. For serum samples, nine concentrations (3.91, 7.81, 15.62, 31.25, 62.5, 125, 250, 500, 1000ng/mL) of an IVM stock standard prepared in methanol were added to chicken serum not treated with IVM. 200ng/mL ABM was added as the internal standard to assess extraction efficiency by comparison of the peak areas from fortified blank samples with the peak area of the added ABM. Standard curves were injected three times into the chromatography system to confirm accuracy and precision. The concentration of analyte present in a sample was then calculated following interpolation from the described calibration curve.

### Statistical analyses

Statistical analyses were performed using GraphPad Prism Version 10.6.0. Survival in mosquito bioassays was analyzed using Kaplan-Meier survival curves and compared using Mantel-Cox (log-rank) tests. Details regarding statistical tests comparing bird feed consumption rates and IVM sera concentrations are described in respective figure legends. IVM sera concentrations from birds were correlated to cumulative mosquito morality from bioassays conducted on respective animals using Spearman correlation.

## Results

To determine which species we could target in developing IVM-treated feed for field evaluation, we identified host blood meals from *Cx. tarsalis* collected from cities and towns located in the South Platte River valley in Northern Colorado during WNV transmission season (**Additional file 1: Table S1**). The majority of *Cx. tarsalis* blood meals came from avian hosts (84%) (**Fig. 1A, Additional file 1: Table S2**). Of these avian blood meals, 33% came from house sparrows, with each other detected avian species representing less than 20% of identified blood meal hosts (Fig. 1B, Additional file 1: Table S2). Mosquitoes fed on house sparrows throughout the WNV transmission season and were detected at 6/13 sample sites (**Fig. 1C, Additional file 1: Table S2**).

**Figure 1.**
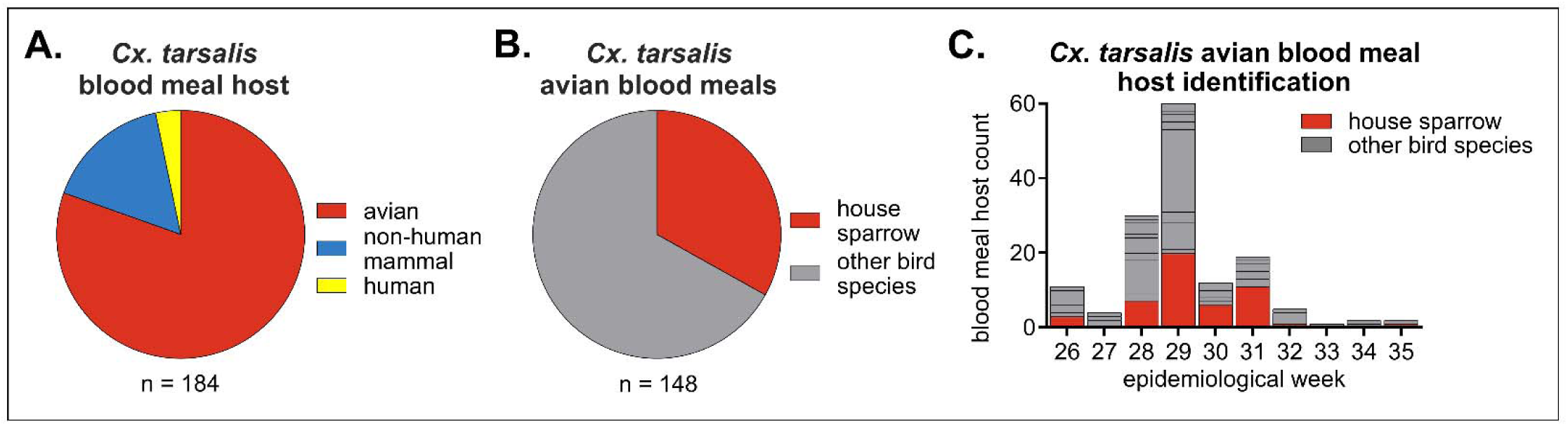
Blood meals identified from *Cx. tarsalis* captured in Northern Colorado indicate house sparrows are a primary target. *Cx. tarsalis* mosquitoes (n = 184) were trapped throughout Northern Colorado in summer 2023. **A)** Blood meal host identification. **B)** Avian blood meal host identification. All species and sample site details are listed in **Additional file 1: Table S2**. **C)** Avian blood meal host identification throughout the WNV transmission season (epidemiological week 26-35).

### Field-collected *Cx. tarsalis* are susceptible to IVM via avian blood meals

We previously determined consumption of 200 mg/kg of IVM-treated feed did not result in any adverse events in chickens and caused their sera to be mosquitocidal to colony-raised *Cx. tarsalis* mosquitoes [29]. To validate the efficacy of this formulation and dose in field-collected mosquitoes, we captured adult *Cx. tarsalis* in Larimer County, CO in July through August 2021 (**Additional file 1: Table S4**). Four-week-old chickens (*Gallus gallus*) were exclusively fed 200 mg/kg powdered IVM in mash for three consecutive weeks. All chickens remained healthy and exhibited no clinical symptoms with all scores reported as “A” per our behavioral protocol, and weight gain was similar between IVM and control feed groups (**Fig. 2A**). Consistent with our previous studies, colonized *Cx. tarsalis* experienced significant mortality after feeding on blood from IVM-treated chickens compared to control, with 80% mortality after seven days post feeding (n = 72 mosquitoes fed on control sera, n = 91 mosquitoes fed on IVM-treated sera, Mantel-Haenszel hazard ratio = 3.614, 95% confidence interval (CI) = 2.300-5.679) (**Fig. 2B**). Similarly, field-collected *Cx. tarsalis* fed on IVM-treated chickens experienced significant mortality, though they had lower mortality overall (42%) (n = 68 mosquitoes fed on control sera, n = 79 mosquitoes fed on IVM-treated sera, Mantel-Haenszel hazard ratio = 3.908, 95% CI = 2.064-7.401) (**Fig. 2C**). Due to the challenge in collecting field mosquitoes and feeding them directly on live birds, we conducted all subsequent bioassays in colonized *Cx. tarsalis* mosquitoes.

**Figure 2.**
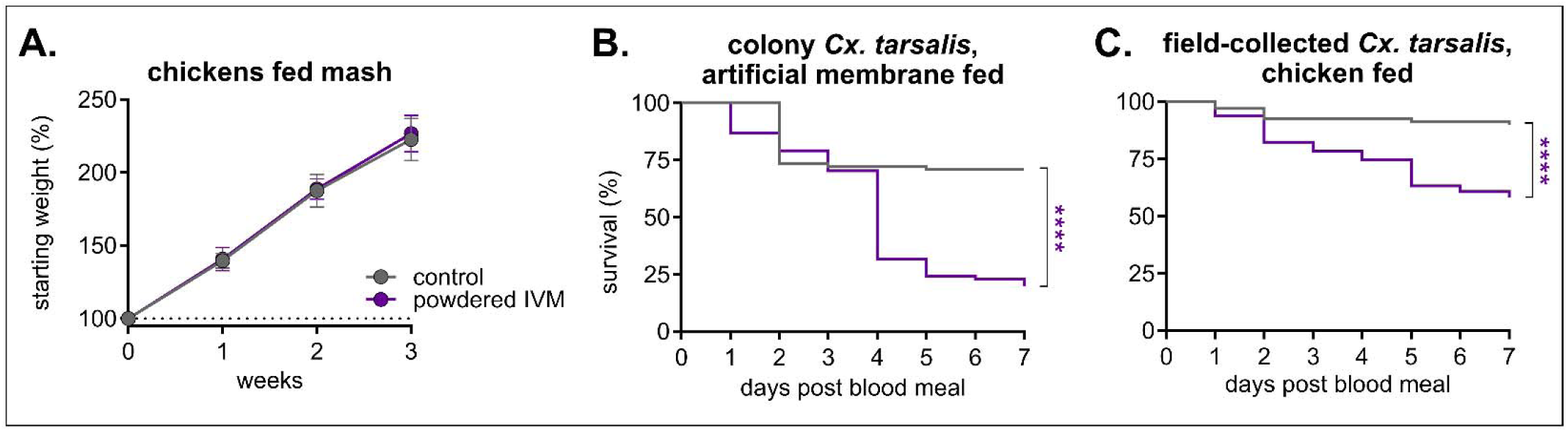
Field-collected *Cx. tarsalis* mosquitoes experience significant mortality when fed chicken blood containing IVM. Chickens (n = 6 in control group, n = 5 in IVM-treated group) were fed control or 200 mg/kg of powdered IVM-treated mash for three weeks. **A)** Chickens were weighed weekly (mean ± SD). **B)** Colony and **(C)** field-collected *Cx. tarsalis* were fed **B)** indirectly via an artificial membrane feeder or **C)** directly on chickens, and mortality was measured over seven days. Log-rank (Mantel-cox) test, ****p<0.0001.

### High-dose IVM-treated feed had a strong safety profile in pigeons and was mosquitocidal to *Cx. tarsalis*

After observing less mortality in field-captured compared to colonized *Cx. tarsalis*, we formulated a high-dose IVM diet (400 mg/kg) using IVM sprayed onto commonly used seed diets (uncoated IVM) to increase mosquitocidal efficacy in a field setting. Sunflower seed and millet were chosen to target a wide range of species likely to local bird feeders [46]. Domestic pigeons (*Columba livia*) were used to evaluate safety and efficacy in a species likely to consume seed from a bird feeder. Pigeons in all groups appeared healthy throughout and after eating the formulated diet, with similar rates of seed consumption (**Fig. 3 A/D**) and weight change (**Fig. 3 B/E**) and clinical scores reported as “A” as per our behavioral protocol. Colonized *Cx. tarsalis* experienced significant mortality after feeding on sera from IVM-treated pigeons compared to control, with 100% mortality after seven days post-blood-feed in both IVM-treated sunflower seed (n = 18 mosquitoes fed on control sera, n = 30 fed on IVM sera, Mantel-Haenszel hazard ratio = 13.8, 95% CI = 5.823-32.71) and millet (n = 11 mosquitoes fed on control sera, n = 10 mosquitoes fed on IVM sera, Mantel-Haenszel hazard ratio = 39.58, 95% CI = 8.81-177.8) groups (**Fig. 3 C/F**).

**Figure 3.**
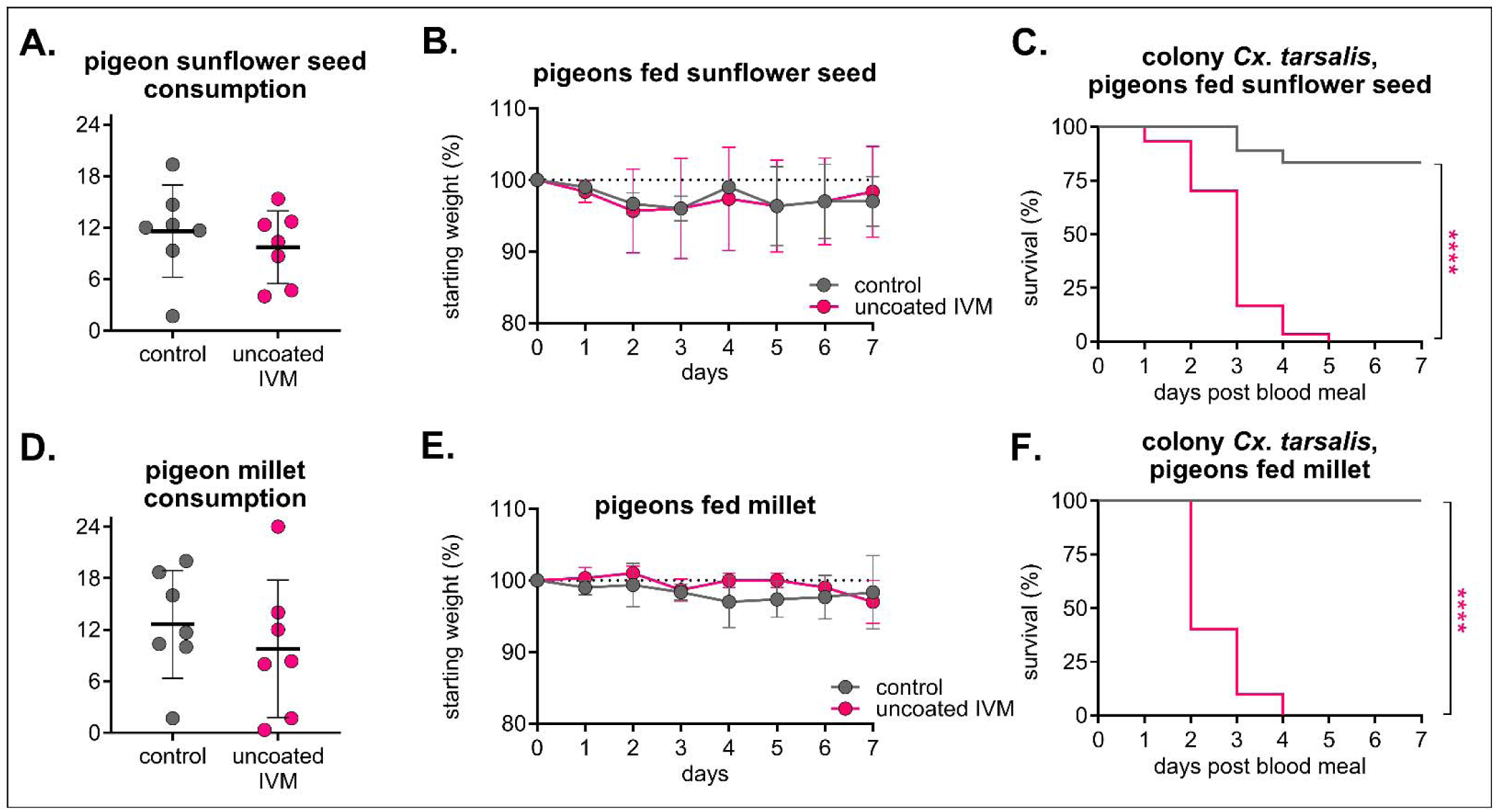
**High-dose IVM-treated sunflower seed and millet were well-tolerated in pigeons and confer significant mortality to *Cx. tarsalis.*** Pigeons (n = 3 per group) were fed either control, or 400 mg/kg uncoated IVM-treated **A-C)** sunflower seed or **D-F)** millet for seven days. **A/D)** Total seed eaten per group was measured daily (mean ± SD). Each point represents seed eaten by group over the course of one day. No significant differences (p>0.05) were observed between groups using an unpaired t-test. **B/E)** Pigeons were weighed daily (mean ± SD). **C/F)** Colony *Cx. tarsalis* were fed pigeon sera collected on day three after feeding on IVM-treated **C)** sunflower seed or **F)** millet via an artificial membrane feeder, and mortality measured over seven days. Log-rank (Mantel-cox) test, ****p<0.0001.

To improve environmental stability and pharmacokinetic properties of the diet, we developed excipient-formulated IVM-treated seeds coated with Opadry® II, a polyvinyl alcohol polymer commonly used in controlled release formulations and tablet coatings. Because we observed complete mortality in mosquitoes fed sera from high-dose IVM-treated birds (**Fig. 3**), we selected the lower-dose formulation for evaluation in following experiments. Pigeons consumed uncoated and coated IVM millet at similar rates compared to control feed (**Fig. 4A**), and all birds were considered healthy throughout the study, with similar weight change in both IVM groups compared to their respective controls and clinical scores reported as “A” as per our behavioral protocol (**Fig. 4B/C**). Additionally, IVM concentrations in pigeon sera were similar within matched timepoints between uncoated and coated IVM groups (**Fig. 4D**). Colonized *Cx. tarsalis* fed day three-collected sera from both IVM-treated groups experienced significant mortality, with over 95% mortality in both IVM groups compared to 97% in control (n = 67 mosquitoes fed on control sera, n = 92 mosquitoes fed on uncoated IVM sera, Mantel-Haenszel hazard ratio = 20.45, 95% CI =12.51-33.44; n = 94 mosquitoes fed on coated IVM sera, Mantel-Haenszel hazard ratio = 23.69, 95% CI 14.45-38.83) (**Fig. 4E**). The same effect was observed in *Cx. tarsalis* fed day seven-collected sera, albeit with comparatively higher survivorship rates of 83% mortality in uncoated and 91% mortality in coated compared to 7% in control (n = 29 mosquitoes fed on control sera, n = 40 mosquitoes fed on uncoated IVM sera, Mantel-Haenszel hazard ratio = 11.77, 95% CI = 5.458-25.37; n = 34 mosquitoes fed on coated IVM sera, Mantel-Haenszel hazard ratio = 18.42, 95% CI = 8.036-42.24) (**Fig. 4F**).

**Figure 4.**
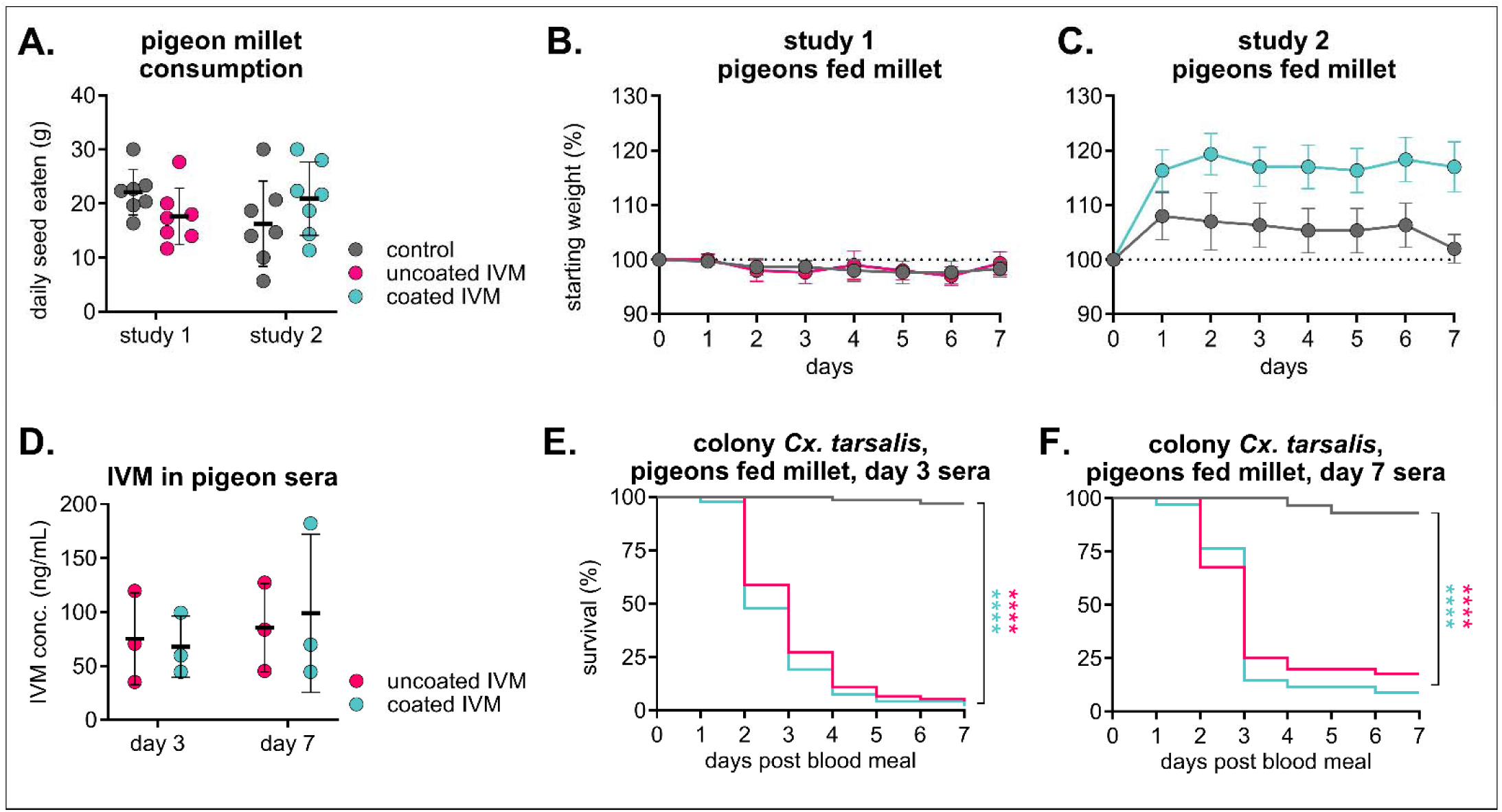
Excipient-formulated millet diets are palatable and efficacious. Pigeons (n = 3 per group) were fed either control or 200mg/kg IVM-treated uncoated (study 1) or Opadry II-coated (study 2) millet for seven days. **A)** Total seed eaten per group was measured daily (mean ± SD). Each point represents seed eaten by group over the course of one day. No significant differences (p>0.05) between groups within study using a two-way ANOVA were observed. **B/C)** Pigeons were weighed daily (mean ± SD). **D)** IVM was measured in pigeon sera and there were no statistical differences within time points (p>0.05) between IVM groups using a two-way ANOVA with with Šídák’s multiple comparisons test. **E/F)** Colony *Cx. tarsalis* were fed pigeon sera collected on E) day three or F) day seven via an artificial membrane feeder, and mortality measured over seven days. Log-rank (Mantel-cox) test, ****p<0.0001.

### Excipient-formulated IVM-treated feed is well-tolerated and effective in small passerine species

We next tested the same formulation (200 mg/kg IVM-treated feed) in zebra finches (*Taeniopygia guttata*) because of their small size, well-documented husbandry and accessibility, establishing it as a valuable animal model to evaluate the approximate effects of this diet on passerine birds [47]. All zebra finches fed 200 mg/kg IVM-treated feed exhibited severe stupor, lethargy, and ataxia within 24 hours of diet provision. After they were checked by the laboratory animal veterinarian, birds were removed from diets and all recovered within 24 hours of withdrawal. After an additional two-week withdrawal period to clear residual IVM, they were given a decreased concentration of 50 mg/kg and monitored three times daily. At this concentration, we did not observe adverse clinical signs throughout the duration of the diet, birds consumed experimental and control seed at similar rates, all scores reported as “A” per our behavioral protocol, and all birds maintained their weight (**Fig. 5A/B**). While still significant compared to mosquitoes fed control sera, we observed less mosquito mortality (61% mortality) in mosquitoes fed sera from coated IVM-treated birds (n = 12 mosquitoes fed on control sera, n = 23 mosquitoes fed on coated IVM sera, Mantel-Haenszel hazard ratio = 8.081, confidence interval 2.380-27.44) (**Fig. 5C**). We then increased the dose to 100 mg/kg IVM, and at this dose, even though birds in the coated IVM millet group consumed more feed than birds in the uncoated IVM millet group (**Fig. 5D**), all zebra finches remained healthy and maintained their weight (**Fig. 5E**). Interestingly, IVM concentrations in sera collected on day seven from zebra finches fed uncoated IVM millet were significantly higher than that of coated IVM millet despite increased consumption in the coated IVM millet group (**Fig. 5F**). Despite these differences, colonized mosquitoes fed sera from both uncoated and coated IVM groups experienced comparable significant mortality, with 100% mortality in uncoated IVM and 96% mortality in coated IVM compared to 27% in control (n = 98 mosquitoes fed on control sera, n = 77 mosquitoes fed on uncoated IVM sera, Mantel-Haenszel hazard ratio = 12.77, confidence interval 7.764-21.01; n = 101 mosquitoes fed on coated IVM sera, Mantel-Haenszel hazard ratio = 8.835, confidence interval 5.667-13.77) (**Fig. 5G**).

**Figure 5.**
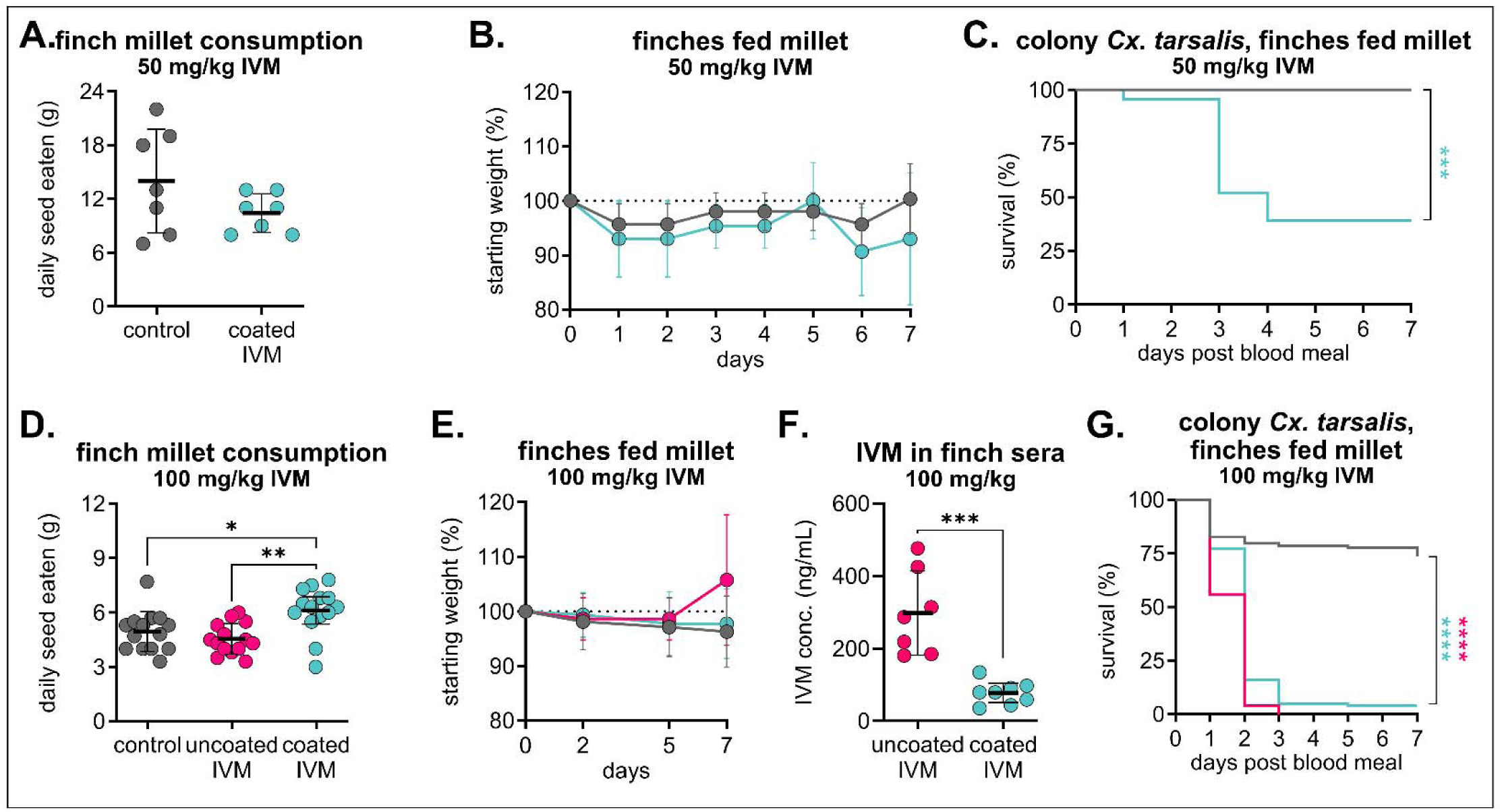
Mosquitocidal efficacy in zebra finch sera varies between doses and excipient formulations. Zebra finches were fed either **A-C)** control or 50 mg/kg IVM-treated Opadry II-coated millet (n = 3 per group) or **D-G)** control or 100 mg/kg IVM-treated uncoated or Opadry II-coated millet (n = 8 per each IVM group, 7 per control group) for seven days. **A/D**) Total seed eaten per group was measured daily (mean ± SD). Each point represents seed eaten by group over the course of one day. **A)** No significant difference (p>0.05) between groups were observed using an unpaired t-test. **D)** One-way ANOVA with Tukey’s multiple comparisons test (*p<0.05, **p<0.005). **B/E)** Zebra finches were weighed **B)** daily or **E)** four times over the seven-day diet period (mean ± SD). **C)** Colony *Cx. tarsalis* were fed zebra finch sera via an artificial membrane feeder, and mortality measured over seven days. Log-rank (Mantel-cox) test, ***p<0.005. **F)** IVM was measured in sera of zebra finches fed 100 mg/kg uncoated and coated IVM millet diets. Concentrations were compared using an unpaired t-test (***p<0.0005) **G)** Colony *Cx. tarsalis* were fed zebra finch sera via an artificial membrane feeder, and mortality measured over seven days. Log-rank (Mantel-cox) test, ****p<0.0001.

After determining that 100 mg/kg IVM feed was safe and efficacious in zebra finches, we evaluated the same dose and formulation in field-caught house sparrows (*Passer domesticus*). We selected house sparrows because they are a primary blood meal source for *Cx. tarsalis* mosquitoes in northern Colorado [41] (**Fig. 1**), and are frequently spotted at bird feeders [46]. All birds remained apparently healthy throughout the diet, and no differences in seed consumption nor weight change were observed (**Fig. 6A/B**). Evaluation of liver histology revealed mild to moderate chronic hepatitis in all birds, including controls, and an overall lack of overt toxic changes in the treated group (**Fig. 6C, Table S5**). Unlike zebra finches (**Fig. 5D**), IVM concentrations in sera from house sparrows were comparable between uncoated and coated IVM millet (**Fig. 6D**). Colonized mosquitoes fed sera from both uncoated and coated IVM groups experienced comparable significant mortality, with 100% mortality in both uncoated and coated IVM groups compared to 37% in the control (n = 57 mosquitoes fed on control sera, n = 160 mosquitoes fed on uncoated IVM sera, Mantel-Haenszel hazard ratio = 6.025, confidence interval 3.811-9.526; n = 165 mosquitoes fed on coated IVM sera, Mantel-Haenszel hazard ratio = 5.334, confidence interval 3.403-8.363) (**Fig. 6E**). A large proportion of mosquitoes fed sera from the control group experienced die-off one day post-blood-feeding, but remaining mosquitoes showed no signs of toxicity compared to mosquitoes fed sera from either IVM group, which were all paralyzed (**Fig. 6F**).

**Figure 6.**
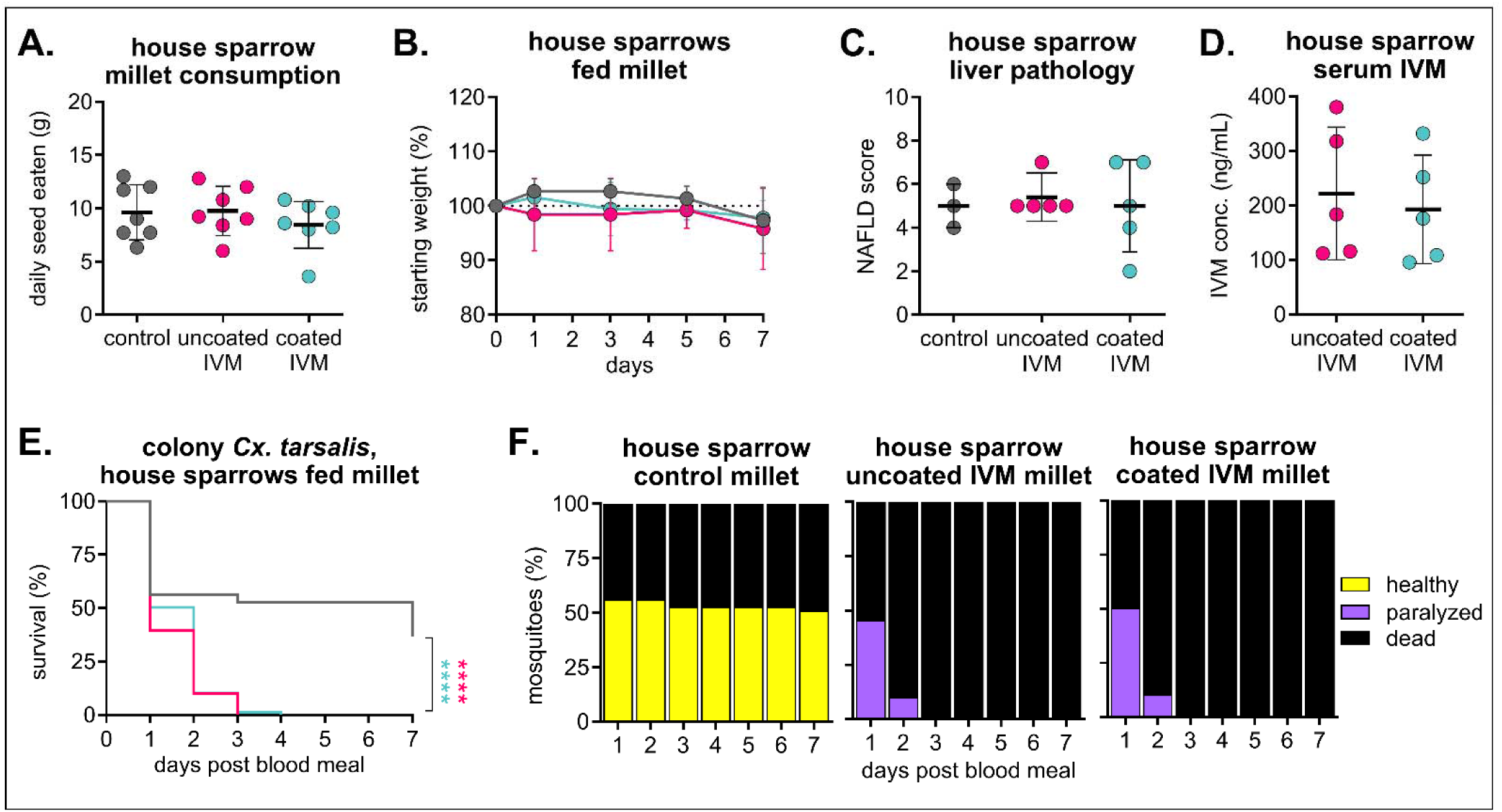
House sparrows safely consume IVM-treated feed and develop mosquitocidal concentrations in sera. House sparrows (n = 3 in control group, n = 5 per each IVM group) were fed 100 mg/kg IVM uncoated or Opadry II-coated millet for seven days. **A)** Total seed eaten per group was measured daily (mean ± SD). No significant difference (p>0.05) between all groups using one-way ANOVA with Tukey’s multiple comparisons test. **B)** House sparrows were weighed every other day (mean ± SD). **C)** House sparrow liver tissue was evaluated and scored according to standards for non-alcoholic fatty liver disease (NAFLD). No significant difference (p>0.05) was observed between groups using one-way ANOVA. **D)** IVM was measured in house sparrow sera collected on day seven of the diet. No significant difference (p>0.05) between groups using unpaired t-test. **E/F)** Colony *Cx. tarsalis* were fed house sparrow sera via an artificial membrane feeder, and **E)** mortality measured over seven days. Log-rank (Mantel-cox) test, ****p<0.0001. **F)** Blood-fed mosquitoes were observed for signs of toxicity for seven days.

### Relationship between IVM formulation, real dose, and mosquitocidal efficacy

To better understand the relationship between IVM formulation and dose, we extrapolated the real dose by calculating the total consumed IVM per bird weight and compared this value to concentration of IVM in diet and IVM concentration in sera from pigeons fed 200 mg/kg formulations, all house sparrows, and all zebra finches (Fig 7A/B, Additional file 1: Table S6). Real dose ranged between 9-40 mg IVM/kg bird weight in all birds that appeared clinically healthy throughout the diet. Zebra finches fed the 200 mg/kg diet, which were the only animals to present signs of IVM toxicity following diet ingestion, received a real dose of 67 mg IVM/kg bird weight, substantially higher than other birds. Further, we correlated IVM concentration in individual birds’ sera to *Cx. tarsalis* survivorship (Fig. 7C, Additional file 1: Table S7). There were no observable relationships between species, formulation, and survivorship (Additional file 1: Table S8). Overall, *Cx. tarsalis* survivorship was below 50% in all groups irrespective of the real doses or IVM concentrations in sera, and at or below 40% in bioassays conducted with serum from passerine birds.

**Figure 7.**
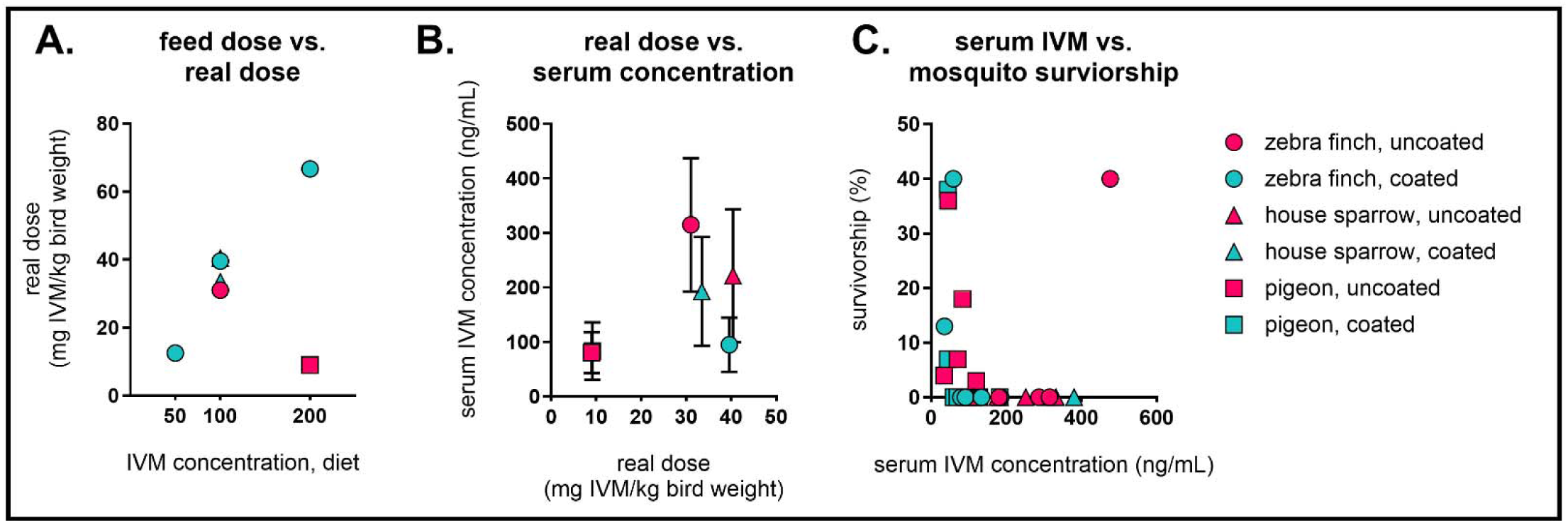
Relationships between IVM formulation, real dose, and mosquitocidal efficacy. Comparisons between real dose, IVM concentration in diet, IVM concentration in sera, and mosquito survivorship. **A)** IVM dose in diet compared to real dose per average weight of birds. Real dose is calculated as follows: (kg of feed eaten per group × mg of IVM in diet)/(average weight of birds in group in kg). **B)** Real dose per average weight of birds compared to IVM concentration in bird sera. **C)** Individual avian IVM serum concentrations versus corresponding cumulative *Cx. tarsalis* survivorship on day seven post blood feeding.

## Discussion

We report here that IVM-treated bird feed is likely to reach the preferred blood meal source of WNV vector *Cx. tarsalis* mosquitoes, has a strong safety profile for many avian species consuming it, and confers mosquitocidal efficacy to treated-bird serum when fed to both colonized and field-collected *Cx. tarsalis*. Birds as small as 11g can tolerate eating exclusively 100mg IVM per kg of bird feed for up to one week with no evidence of adverse clinical signs nor histopathology. Colonized *Cx. tarsalis* are susceptible to serum from passerine birds fed formulated diets at 100 mg/kg, suggesting that this strategy can be effective at reducing WNV transmission in the field by targeting blood-feeding mosquitoes that prefer to feed on birds. Together, these data indicate that IVM-treated bird feed may be an effective WNV transmission control strategy via targeting blood feeding adult mosquitoes when implemented in areas of high WNV transmission between songbirds and ornithophilic mosquitoes.

*Cx. tarsalis* feeding behavior, as identified by blood meal analysis presented in this study, largely corroborated findings from previously published studies conducted in Colorado and other western states [41,48], with the majority of blood meals derived from avian hosts. A large proportion of blood meals came from common visitors to bird feeders, including house sparrows, mourning doves, house finches, and black-capped chickadees. We previously assessed spatial movement of common birdfeeder species relative to bird feeder sites to determine how optimal deployment of IVM-treated feeders could reduce local transmission throughout the WNV season by reducing enzootic transmission prior to the onset of human infections [31]. In that study, we tagged house finches in Fort Collins, CO, with passive integrated transponders to quantify their visits to bird feeders [31]. Of the 2,082 bird feeder visits, the average duration was ∼21 seconds and an individual bird was detected visiting a feeder 19 times per day. These data suggest small passerines will frequent the feeders often each day in the summer, coinciding with peak *Cx. tarsalis* activity. The success of an IVM-treated bird feed strategy for reducing virus transmission depends on *Cx. tarsalis* taking blood meals from self-medicated birds, so these data together with our blood meal host identification results suggest that mosquitoes are likely to take blood meals from house sparrows and other passerine birds that frequently visit bird feeders. All mosquitoes in this study were collected in CDC light traps baited with CO_2_, which attract host-seeking female mosquitoes. This dataset does not include mosquitoes captured in resting and gravid traps, which could bias blood meal distributions [49]. However, it is well-documented that *Cx. tarsalis* primarily feed on birds and opportunistically feed on mammals, and our findings show the same overall distribution.

We developed IVM-seed formulations that are attractive to a broad variety of birds and can modulate drug release upon ingestion. Using both hulled millet and hulled sunflower seeds enables preferential targeting of both small and medium sized granivorous birds, and we found comparable rates of consumption between treated and untreated seed within seed type and species groups. For IVM to be successful in the field, it also needs to be stable on a feed product. We also evaluated IVM-seed formulations coated with aqueous excipient Opadry® II to maintain the stability of the active ingredient on the seeds and modulate controlled release of the drug in the avian digestive system [50]. Most birds consumed both uncoated and coated IVM feeds at similar rates to control, though zebra finches consumed more coated IVM feed than either uncoated or control. Interestingly, this did not result in an increased IVM concentration in sera – rather, zebra finches fed coated IVM exhibited lower concentrations than those fed uncoated IVM. This could be attributed to Opadry® II suppressing the maximum concentration (C_max_), delaying C_max_ to peak later over the course of exposure, and extending the period of bioavailability. Conventionally, therapeutic doses of IVM in companion birds, poultry, and captive raptors range from 0.2-2.0 mg IVM/kg body weight [51,52], with severe toxicity observed in chickens at 15 mg IVM/kg body weight delivered subcutaneously [53]. IVM selectively binds to invertebrate glutamate-gated chloride channels, but at high enough concentrations, it can potentiate the structurally similar vertebrate γ-aminobutyric acid (GABA_A_)-gated chloride channels that inhibit interneurons in the central nervous system (CNS). To access these (GABA_A_)-gated chloride channels, IVM could pass beyond the blood brain barrier by oversaturating P-glycoprotein transporters that limit drug uptake into the brain [54]. To prevent reaching this limit of tolerance, future studies should characterize pharmacokinetic profiles of IVM metabolism over time and throughout different tissues in a variety of avian species. Overall, even though our extrapolated real doses were higher than previously documented toxic doses, IVM treated birdseed was safe up to 100mg/kg coated feed, with no clinical nor histological signs of toxicity in any of the birds evaluated at this concentration. Importantly, in our experiments, birds ate only IVM-treated feed for seven consecutive days, whereas in field trials, we would expect wild birds to supplement their diet with non-treated feed. Therefore, since we did not observe adverse effects in our laboratory birds at the target dose of 100 mg/kg, we do not expect to observe them in the field.

Both colonized and field-collected *Cx. tarsalis* were susceptible to sera from IVM-treated birds at all diet concentrations, however field-collected mosquitoes had higher rates of survival. Interestingly, we saw no relationship between mosquitocidal effect and type of seed (e.g., millet, sunflower seed), IVM formulation (e.g., coated, uncoated), or sera IVM concentration demonstrating that excipient does not inhibit the ability of the treated bird’s sera from affecting mosquito survival. We do not expect field-collected *Cx. tarsalis* to have developed targeted resistance to IVM, and their higher survival may be due to increased fitness of wild mosquitoes compared to highly inbred and genetically restricted laboratory mosquitoes such as ours which were colonized in the 1960s [55]. Because the majority of data presented in this study were generated from studies conducted with colonized mosquitoes, future experiments should determine survivorship in genetically diverse, field-collected mosquitoes to evaluate susceptibility in wild mosquito populations. Studies in other mosquito genera including *Culex* and *Anopheles* have specifically implicated ATP binding cassette transporters and cytochrome P450 monooxygenases (CYP P450) in IVM metabolism [56–58]. Mutations in several different detoxification enzymes including CYP P450, hydrolases, and glutathione S-transferases (GST) have been identified in field-collected insecticide-resistant populations [59], increasing the likelihood that field-collected *Cx. tarsalis* may have a broad assortment of alleles that could contribute to functional resistance. However, we observed that both field-collected and colonized *Cx. tarsalis* typically died within the first three days after exposure to IVM in avian blood meals, and most IVM-exposed mosquitoes exhibited some degree of flaccid paralysis before recovery. These mosquitoes would be expected to die under typical competitive field conditions, suggesting increased efficacy compared to what we observed in field-collected *Cx. tarsalis* under laboratory conditions. Overall, we observed mosquitocidal efficacy in *Cx. tarsalis* indirectly and directly fed sera from birds fed IVM-treated feed, with no correlation between real dose, IVM concentration, and mosquito survivorship. *In vivo* blood-feeding data show that avian IVM concentrations needed to kill most *Cx. tarsalis* are significantly less than what we measured *in vitro* serum-replacement membrane feeding experiments (LC_50_ 49.94 ng/mL) [30].

## Conclusions

The results of this study show that 100 mg/kg IVM-treated bird feed is safe for consumption by small passerines and mosquitocidal to blood feeding *Culex tarsalis*. Because we aim to achieve the minimum effective dose with the lowest probability of adverse events in wild birds, we plan to move forward with 100 mg/kg in future trials to evaluate the impact of IVM-treated feed on reducing WNV transmission in the field.

## Abbreviations

## Acknowledgements

The authors would like to thank Nurul Islam, Paul Soma, and John Belisle for assistance with LCMS method development and instrumentation. We thank Angela Bosco-Lauth for feral pigeon capture and training on animal techniques. We thank Allison Vilander for training on sample preparation for histopathological analysis. We thank Talia Wong and Rebekah Kading for their help with optimizing the blood meal identification protocol. We thank Ali Brehm and Tim Burton for assistance with animal sample collections. We are grateful to the volunteer households that allowed us to collect mosquitoes in summer 2023. Graphical abstract was created in BioRender. Ebel, G. (2025) https://BioRender.com/lizwjdp

## Author contributions

MJS: Conceptualization, data curation, formal analysis, investigation, methodology, project administration, validation, visualization, writing – original draft, writing – review and editing; KC: Data curation, formal analysis, investigation, methodology; CMS: Data curation, formal analysis, investigation, methodology, validation, writing – review and editing; CN: Data curation, formal analysis, investigation, methodology, supervision, validation; MER: Investigation; AL: Investigation; PS: Investigation, TP: Investigation; CP: Data curation, investigation; JB: Data curation, formal analysis; JR: Investigation, writing – review and editing, PL: Investigation, supervision; ENG: Formal analysis, visualization, writing – review and editing; BC: Funding acquisition, resources; CB: Funding acquisition; GDE: Funding acquisition, resources, supervision; BDF: Conceptualization, data curation, formal analysis, funding acquisition, investigation, methodology, project administration, resources, supervision, validation, writing – original draft, writing – review and editing.

## Availability of data and materials

Data supporting the conclusions of the present study are included within the article. Data used and/or analyzed during this study are available from the corresponding author upon reasonable request.

## Declarations

### Ethics approval and consent to participate

Animal research was completed under CSU IACUC study protocols #1002 and #4643 “Endectocide in birds to control for arbovirus transmission”. Animals were euthanized by intravenous injection with sodium pentobarbital or isoflurane overdose as approved in the IACUC study protocol. Wild house sparrow collection was completed under Colorado Parks and Wildlife Scientific Collection License #2845076549.

### Consent for publication

All authors have reviewed and approved this manuscript for submission.

## Works Cited

1. Roehrig JT. West Nile virus in the United States - A historical perspective. Viruses. 2013;5:3088–108. 10.3390/v5123088

2. Chancey C, Grinev A, Volkova E, Rios M. The global ecology and epidemiology of West Nile virus. BioMed Research International. Hindawi Limited; 2015. 10.1155/2015/376230

3. George TL, Harrigan RJ, Lamanna JA, Desante DF, Saracco JF, Smith TB. Persistent impacts of West Nile virus on North American bird populations. Proceedings of the National Academy of Sciences of the United States of America. 2015; 10.1073/pnas.1507747112

4. U.S. Centers for Disease Control and Prevention. West Nile Virus Historic Data (1999-2024). Accessed 23 Oct 2025.

5. Bolling BG, Moore CG, Anderson SL, Blair CD, Beaty BJ. Entomological studies along the Colorado front range during a period of intense West Nile virus activity. Journal of the American Mosquito Control Association. 2007;23:37–46. 10.2987/8756-971X(2007)23[37:ESATCF]2.0.CO;2

6. Fauver JR, Pecher L, Schurich JA, Bolling BG, Calhoon M, Grubaugh ND, et al. Temporal and spatial variability of entomological risk indices for West Nile virus infection in Northern Colorado: 2006-2013. Journal of Medical Entomology. 2016;53:425–34. 10.1093/jme/tjv234

7. Ronca SE, Murray KO, Nolan MS. Cumulative incidence of West Nile Virus infection, continental United States, 1999–2016. Emerging Infectious Diseases. Centers for Disease Control and Prevention (CDC); 2019;25:325–7. 10.3201/eid2502.180765

8. Sejvar JJ. The long-term outcomes of human West Nile virus infection. Clinical Infectious Diseases. 2007. p. 1617–24. 10.1086/518281

9. Staples JE, Shankar MB, Sejvar JJ, Meltzer MI, Fischer M. Initial and long-term costs of patients hospitalized with West Nile virus disease. American Journal of Tropical Medicine and Hygiene. 2014; 10.4269/ajtmh.13-0206

10. Naveed A, Eertink LG, Wang D, Li F. Lessons Learned from West Nile Virus Infection: Vaccinations in Equines and Their Implications for One Health Approaches. Viruses. Switzerland; 2024;16. 10.3390/v16050781

11. Gould CV, Staples JE, Huang CY-H, Brault AC, Nett RJ. Combating West Nile Virus Disease — Time to Revisit Vaccination. New England Journal of Medicine. Massachusetts Medical Society; 2023;388:1633–6. 10.1056/nejmp2301816

12. Fox JL. Uncertainties in Tracking West Nile Virus Undermine Vaccine Development. Microbe Magazine. 2015; 10.1128/microbe.10.98.1

13. Dawson D, Salice CJ, Subbiah S. The Efficacy of the *Bacillus thuringiensis israelensis* Larvicide Against *Culex tarsalis* in Municipal Wastewater and Water from Natural Wetlands. Journal of the American Mosquito Control Association. 2019;35:97–106. 10.2987/18-6771.1

14. Ruktanonchai DJ, Stonecipher S, Lindsey N, McAllister J, Pillai SK, Horiuchi K, et al. Effect of aerial insecticide spraying on West Nile virus disease-north-central Texas, 2012. American Journal of Tropical Medicine and Hygiene. 2014;91:240–5. 10.4269/ajtmh.14-0072

15. Rhyne MN, Richards SL. Impact of the Insect Growth Regulator Pyriproxyfen on Immature Development, Fecundity, and Fertility of *Aedes albopictus*. Journal of the American Mosquito Control Association. 2020;36:11–5. 10.2987/19-6893.1

16. Thier A. Balancing the risks: vector control and pesticide use in response to emerging illness. Journal of Urban Health: Bulletin of the New York Academy of Medicine. 2001;78:372–81. 10.1093/jurban/78.2.372

17. Darboux I, Charles JF, Pauchet Y, Warot S, Pauron D. Transposon-mediated resistance to Bacillus sphaericus in a field-evolved population of *Culex pipiens* (Diptera: Culicidae). Cellular Microbiology. 2007;9:2022–9. 10.1111/j.1462-5822.2007.00934.x

18. Su T, Thieme J, White GS, Lura T, Mayerle N, Faraji A, et al. High Resistance to Bacillus sphaericus and Susceptibility to Other Common Pesticides in *Culex pipiens* (Diptera: Culicidae) from Salt Lake City, UT. Journal of Medical Entomology. Oxford University Press; 2019;56:506–13. 10.1093/jme/tjy193

19. Holcomb KM, Reiner RC, Barker CM. Spatio-temporal impacts of aerial adulticide applications on populations of West Nile virus vector mosquitoes. Parasites Vectors. 2021;14:120. 10.1186/s13071-021-04616-6

20. Macedo PA, Schleier JJ 3rd, Reed M, Kelley K, Goodman GW, Brown DA, et al. Evaluation of efficacy and human health risk of aerial ultra-low volume applications of pyrethrins and piperonyl butoxide for adult mosquito management in response to West Nile virus activity in Sacramento County, California. J Am Mosq Control Assoc. United States; 2010;26:57–66. 10.2987/09-5961.1

21. Barber LM, Schleier JJ, Peterson RKD. Economic cost analysis of West Nile virus outbreak, Sacramento County, California, USA, 2005. Emerging Infectious Diseases. 2010;16:480–6. 10.3201/eid1603.090667

22. Huff Hartz KE, Weston DP, Johanif N, Poynton HC, Connon RE, Lydy MJ. Pyrethroid bioaccumulation in field-collected insecticide-resistant Hyalella azteca. Ecotoxicology. 2021;30:514–23. 10.1007/s10646-021-02361-1

23. Crowder J, Rochlin I, Bibbs CS, Pennock E, Browning M, Lott C, et al. Managed honey bees, Apis mellifera (Hymenoptera: Apidae), face greater risk from parasites and pathogens than mosquito control insecticide applications. Sci Total Environ. 2025;964:178638. 10.1016/j.scitotenv.2025.178638

24. Oberhauser KS, Manweiler SA, Lelich R, Blank M, Batalden RV, Anda AD. Impacts of ultra-low volume resmethrin applications on non-target insects. Journal of the American Mosquito Control Association. 2009;25:83–93. 10.2987/08-5788.1

25. Tedesco C, Ruiz M, McLafferty S. Mosquito politics: Local vector control policies and the spread of West Nile virus in the Chicago region. Health & Place. 2010;16:1188–95. 10.1016/j.healthplace.2010.08.003

26. Paull SH, Horton DE, Ashfaq M, Rastogi D, Kramer LD, Diffenbaugh NS, et al. Drought and immunity determine the intensity of West Nile virus epidemics and climate change impacts. Proc R Soc B. 2017;284:20162078. 10.1098/rspb.2016.2078

27. Hamer GL, Kitron UD, Goldberg TL, Brawn JD, Loss SR, Ruiz MO, et al. Host Selection by *Culex pipiens* Mosquitoes and West Nile Virus Amplification. Am J Trop Med Hyg. 2009;80:268–78. 10.4269/ajtmh.2009.80.268

28. Brinton M. Replication Cycle and Molecular Biology of the West Nile Virus. Viruses. 2013;6:13–53. 10.3390/v6010013

29. Nguyen C, Gray M, Burton TA, Foy SL, Foster JR, Gendernalik AL, et al. Evaluation of a novel West Nile virus transmission control strategy that targets *Culex tarsalis* with endectocide-containing blood meals. Armstrong PM, editor. PLoS Negl Trop Dis. 2019;13:e0007210. 10.1371/journal.pntd.0007210

30. Holcomb KM, Nguyen C, Foy BD, Ahn M, Cramer K, Lonstrup ET, et al. Effects of ivermectin treatment of backyard chickens on mosquito dynamics and West Nile virus transmission. PLoS Neglected Tropical Diseases. Public Library of Science; 2022;16. 10.1371/journal.pntd.0010260

31. Holcomb KM, Nguyen C, Komar N, Foy BD, Panella NA, Baskett ML, et al. Predicted reduction in transmission from deployment of ivermectin-treated birdfeeders for local control of West Nile virus. Epidemics. Elsevier B.V.; 2023;44. 10.1016/j.epidem.2023.100697

32. Canga AG, Prieto AMS, Liébana MJD, Martínez NF, Vega MS, Vieitez JJG. The pharmacokinetics and metabolism of ivermectin in domestic animal species. Veterinary Journal. 2009;179:25–37. 10.1016/j.tvjl.2007.07.011

33. Martin RJ, Robertson AP, Choudhary S. Ivermectin: An Anthelmintic, an Insecticide, and Much More. Trends in Parasitology. 2021;37:48–64. 10.1016/j.pt.2020.10.005

34. Tesh RB, Guzman H. Mortality and infertility in adult mosquitoes after the ingestion of blood containing ivermectin. Am J Trop Med Hyg. United States; 1990;43:229–33. 10.4269/ajtmh.1990.43.229

35. Meyers JI, Gray M, Kuklinski W, Johnson LB, Snow CD, Black WC 4th, et al. Characterization of the target of ivermectin, the glutamate-gated chloride channel, from *Anopheles gambiae*. J Exp Biol. England; 2015;218:1478–86. 10.1242/jeb.118570

36. Foy BD, Alout H, Seaman JA, Rao S, Magalhaes T, Wade M, et al. Efficacy and risk of harms of repeat ivermectin mass drug administrations for control of malaria (RIMDAMAL): a cluster-randomised trial. Lancet. England; 2019;393:1517–26. 10.1016/S0140-6736(18)32321-3

37. Somé AF, Somé A, Sougué E, Ouédraogo COW, Da O, Dah SR, et al. Safety and efficacy of repeat ivermectin mass drug administrations for malaria control (RIMDAMAL II): a phase 3, double-blind, placebo-controlled, cluster-randomised, parallel-group trial. Lancet Infect Dis. United States; 2025;25:737–50. 10.1016/S1473-3099(24)00751-5

38. Pooda SH, Mouline K, De Meeûs T, Bengaly Z, Solano P. Decrease in survival and fecundity of *Glossina palpalis gambiensis vanderplank* 1949 (Diptera: Glossinidae) fed on cattle treated with single doses of ivermectin. Parasit Vectors. England; 2013;6:165. 10.1186/1756-3305-6-165

39. Borchert JN, Enscore RE, Eisen RJ, Atiku LA, Owor N, Acayo S, et al. Evaluation of rodent bait containing imidacloprid for the control of fleas on commensal rodents in a plague-endemic region of northwest Uganda. J Med Entomol. England; 2010;47:842–50. 10.1603/me09221

40. Mayton EH, Tramonte AR, Wearing HJ, Christofferson RC. Age-structured vectorial capacity reveals timing, not magnitude of within-mosquito dynamics is critical for arbovirus fitness assessment. Parasit Vectors. England; 2020;13:310. 10.1186/s13071-020-04181-4

41. Kent R, Juliusson L, Weissmann M, Evans S, Komar N. Seasonal blood-feeding behavior of *Culex tarsalis* (Diptera: culicidae) in weld County, Colorado, 2007. Journal of Medical Entomology. 2009;46:380–90. 10.1603/033.046.0226

42. Ivanova NV, Zemlak TS, Hanner RH, Hebert PDN. Universal primer cocktails for fish DNA barcoding. Molecular Ecology Notes. John Wiley & Sons, Ltd; 2007;7:544–8. 10.1111/j.1471-8286.2007.01748.x

43. Cicero C, Johnson NK. Higher-Level Phylogeny of New World Vireos (Aves: Vireonidae) Based on Sequences of Multiple Mitochondrial DNA Genes. Molecular Phylogenetics and Evolution. 2001;20:27–40. 10.1006/mpev.2001.0944

44. Meece JK, Reynolds CE, Stockwell PJ, Jenson TA, Christensen JE, Reed KD. Identification of mosquito bloodmeal source by terminal restriction fragment length polymorphism profile analysis of the cytochrome B gene. J Med Entomol. England; 2005;42:657–67. 10.1093/jmedent/42.4.657

45. Bousema T, Dinglasan RR, Morlais I, Gouagna LC, Warmerdam T van, Awono-Ambene PH, et al. Mosquito feeding assays to determine the infectiousness of naturally infected Plasmodium falciparum gametocyte carriers. PLoS ONE. 2012;7. 10.1371/journal.pone.0042821

46. Horn DJ, Johansen SM, Wilcoxen TE. Seed and feeder use by birds in the United States and Canada. Wildlife Society Bulletin. Alliance Communications Group; 2014;38:18–25. 10.1002/wsb.365

47. Patterson MM, Fee MS. Chapter 23 - Zebra Finches in Biomedical Research. In: Fox JG, Anderson LC, Otto GM, Pritchett-Corning KR, Whary MT, editors. Laboratory Animal Medicine (Third Edition) [Internet]. Boston: Academic Press; 2015. p. 1109–34. 10.1016/B978-0-12-409527-4.00023-7

48. Thiemann TC, Lemenager DA, Kluh S, Carroll BD, Lothrop HD, Reisen WK. Spatial Variation in Host Feeding Patterns of *Culex tarsalis* and the *Culex pipiens* complex (Diptera: Culicidae) in California. Journal of Medical Entomology. 2012;49:903–16. 10.1603/ME11272

49. Thiemann TC, Reisen WK. Evaluating sampling method bias in *Culex tarsalis* and *Culex quinquefasciatus* (Diptera: Culicidae) bloodmeal identification studies. J Med Entomol. England; 2012;49:143–9. 10.1603/me11134

50. Koo OMY, Fiske JD, Yang H, Nikfar F, Thakur A, Scheer B, et al. Investigation into Stability of Poly(Vinyl Alcohol)-Based Opadry® II Films. AAPS PharmSciTech. 2011;12:746–54. 10.1208/s12249-011-9630-1

51. Lierz M. Evaluation of the dosage of ivermectin in falcons. Veterinary Record. 2001;148:596–600. 10.1136/vr.148.19.596

52. Hoppes SM. Parasitic Diseases of Pet Birds [Internet]. MSD Veterinary Manual; 2021. https://www.msdvetmanual.com/exotic-and-laboratory-animals/pet-birds/parasitic-diseases-of-pet-birds

53. Kim JS, Crichlow EC. Clinical signs of ivermectin toxicity and the efficacy of antigabaergic convulsants as antidotes for ivermectin poisoning in epileptic chickens. Vet Hum Toxicol. United States; 1995;37:122–6.

54. Scott NE. Chapter 93 - Ivermectin Toxicity. In: Silverstein DC, Hopper K, editors. Small Animal Critical Care Medicine [Internet]. Saint Louis: W.B. Saunders; 2009. p. 392–4. 10.1016/B978-1-4160-2591-7.10093-1

55. Tempelis CH, Reeves WC, Bellamy RE, Lofy MF. A Three-Year Study of the Feeding Habits of Culex Tarsalis in Kern County, California. The American Journal of Tropical Medicine and Hygiene. 1965;14:170–7. 10.4269/ajtmh.1965.14.170

56. Chaccour CJ, Hammann F, Alustiza M, Castejon S, Tarimo BB, Abizanda G, et al. Cytochrome P450/ABC transporter inhibition simultaneously enhances ivermectin pharmacokinetics in the mammal host and pharmacodynamics in *Anopheles gambiae*. Scientific Reports. 2017;7:1–12. 10.1038/s41598-017-08906-x

57. Nicolas P, Kiuru C, Wagah MG, Muturi M, Duthaler U, Hammann F, et al. Potential metabolic resistance mechanisms to ivermectin in *Anopheles gambiae*: a synergist bioassay study. Parasites & Vectors. 2021;14:172. 10.1186/s13071-021-04675-9

58. Buss DS, Mccaffery AR, Callaghan A. Evidence for p-glycoprotein modification of insecticide toxicity in mosquitoes of the *Culex pipiens* complex. Medical and Veterinary Entomology. John Wiley & Sons, Ltd; 2002;16:218–22. 10.1046/j.1365-2915.2002.00365.x

59. Xu Q, Liu H, Zhang L, Liu N. Resistance in the mosquito, *Culex quinquefasciatus*, and possible mechanisms for resistance. Pest Management Science. John Wiley & Sons, Ltd; 2005;61:1096–102. 10.1002/ps.1090

60. Wehmeyer ML, Sauer FG, Lühken R. A minimum data standard for reporting host-feeding patterns of vectors [Internet]. In Review; 2024 [cited 2025 Oct 14]. 10.21203/rs.3.rs-3896902/v1

